# The landscape of interactions between cancer polygenic risk scores and somatic alterations in cancer cells

**DOI:** 10.1101/2020.09.28.316851

**Authors:** Eduard Porta-Pardo, Rosalyn Sayaman, Elad Ziv, Alfonso Valencia

## Abstract

Over the last 15 years we have identified hundreds of inherited variants that increase the risk of developing cancer. Polygenic risk scores (PRS) summarize the genetic risk of each individual by accounting for the unique combination of risk alleles in their genome. So far, most studies of PRS in cancer have focused on their predictive value: i.e. to what extent the PRS can predict which individuals will develop a particular cancer type. In parallel, for most cancers, we have identified several subtypes based on their somatic molecular properties. However, little is known about the relationship between the somatic molecular subtypes of cancer and PRS and it is possible that PRS preferentially predict specific cancer subtypes. Since cancer subtypes can have very different outcomes, treatment options and molecular vulnerabilities, answering this question is very important to understand the consequences that widespread PRS use would have in which tumors are detected early.

Here we used data from The Cancer Genome Atlas (TCGA) to study the correlation between PRS for different forms of cancer and the landscape of somatic alterations in the tumors developed by each patient. We first validated the predictive power of 8 different PRS in TCGA and describe how PRS for some cancer types are associated with specific molecular subtypes or somatic cancer driver events. Our results highlight important questions that could improve the predictive power of PRS and that need to be answered before their widespread clinical implementation.

## Introduction

Over the last fifteen years the biomedical scientific community has collectively done many genome wide association studies (GWAS) to identify hundreds of germline polymorphisms associated different types of cancer. Although variable across cancer types, the variants discovered so far explain a relatively large fraction of the heritability of many forms of the disease. For example, the 170 variants associated with breast cancer^1^ or the 147 variants associated with prostate cancer^2^ explain 40% and 28%, respectively, of these diseases’ heritability.

Given the complex genetic architecture of most forms of cancer, which usually involve dozens or hundreds of loci, polygenic risk scores (PRS) have emerged as an attractive way to integrate the information of all these variants for each individual to summarize their risk for different cancer types^3^. There are now PRS for breast^4–6^, prostate^2^, lung^7^ or thyroid^8^ cancers among many others. In general, PRS are used to predict cancer risk, and beginning to be tested in clinical settings. The goal is to adjust individual conducts accordingly and, in some cases, assess the need to adapt screening strategies^9^.

In parallel, the study of the relations between genetic predisposition and cancer risk has crystalized in multiple lines of evidence showing interactions between germline variants and tumor associated somatic mutations^10–14^. A particularly interesting aspect is the potential collaboration of the germline variants with cancer driver somatic mutations in cancer activation, an oncogenic mechanism which remain poorly understood and that could help us understand, for example, why some individuals develop cancer despite having only a few, or even non-detectable somatic driver mutations. Given the interplay between germline and somatic variants, it is possible that PGRS for some cancer types preferentially identify specific cancer subtypes, like for example in breast cancer: higher PGRS correlate with estrogen receptor positive tumors^5^.

Here we used The Cancer Genome Atlas^15^ (TCGA) to systematically study the relation between PRS and the somatic landscape of the tumors they predict. We first tested the capacity of current PRS to correctly predict TCGA cancer types and molecular subtypes. For the significant PRS we then tested the correlation with the specific cancer driver mutations. Our results show that there is indeed a strong association between PRS and certain cancer subtypes and somatic mutations. We further elaborate on the important implications of these observations for both, the study of the interplay of germline and somatic variants as oncogenic mechanisms as well as for the implementation of PRS for specific cancer types in the clinic.

## Results

### Polygenic risk scores as predictors of TCGA cancer types

We used data from the GWAS catalog^16^ and the recently published Oncoarray studies^17^ to create 32 different PGRS for 19 different cancer types. In brief, to create the PGRS from the Oncoarray studies we clumped all the loci that were genome-wide statistically significant (p < 5 e-8) in each cohort (glioma^18^, lung^19^, breast^1^, colorectal^20^, ovarian^21^ or prostate cancer^2^) to find the variant in each locus with the strongest signal. Then, we assigned a weight to each risk variant equivalent to log of the odds ratio observed in the corresponding study. In the case of PGRS derived from the GWAS catalog, we only kept the lead genome-wide significant variants (p < 5e-8) identified in studies focused on European ancestry with more than 2000 individuals. We assigned a weight equivalent to the log of the odds ratio for the reported risk allele. We then used the germline genotypes of TCGA samples derived from SNParray data to quantify each PRS in each TCGA individual (**Methods**). As expected, PRS that predict the same or similar cancer types correlate with each other across the TCGA cohorts (**Figure S1**). In the end, we selected the best PRS for each of the 8 cancer types where at least one PRS had enough predictive power (**Methods**).

One of the concerns about any PRS is that it might not be predictive in cohorts beyond the one where it was developed. Moreover, most PRS are developed using cohorts that only have cases and healthy controls. Therefore, the predictive power of the different PRS in TCGA, a new dataset that consists only of patients that suffered different cancer types, remains unknown.

To identify the PRS that predict the correct cancer type in TCGA, we first calculated the fold-enrichment in cases across individuals at the extreme deciles of the PRS when compared to those in the intermediate range of the PRS (**Figure 1, S2 and S3**), as well as their area under the ROC curve (AUC). Out of the 27 different PRS, 14 had an AUC above 0.55, a significant enrichment of cases in the top quintile compared to the bottom one (two-sided Fisher’s test FDR < 0.001) and showed a distribution consistent with the correct prediction of the targeted cancer type: depletion of cases at the bottom of the PRS and enrichment of cases at the top (**Figure 1**).

**Figure 1.**
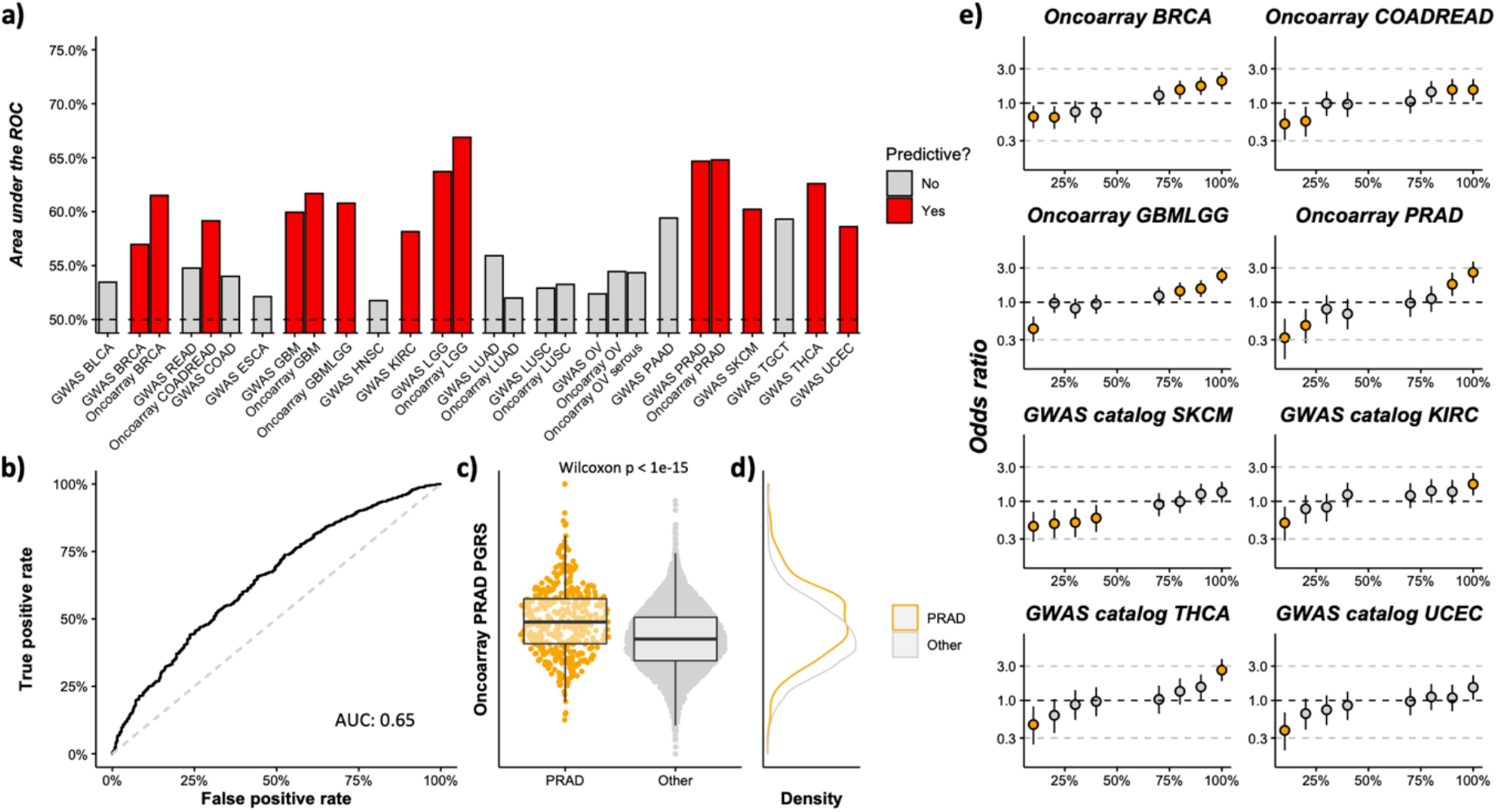
Performance of the PRS in The Cancer Genome Atlas. **a)** We calculated the area under the ROC curve (y-axis) for all the cancer PRS that we create (x-axis). We classified each PRS into predictive (red bars) and not predictive (grey bars) depending on whether they had an AUC > 0.55 and a significant enrichment in cases between the top and bottom quintiles (FRD < 0.001) or not. **b)** ROC curve for the Oncoarray PRAD PRS. **c)** Boxplots showing the distribution of the Oncoarray PRAD PRS values for the TCGA patients we studied. Each dot is a patient and they are grouped according to whether they have had prostate cancer (orange, left) or not (gray, right). **d)** Density curves for the Oncoarray PRAD PRS, following the same color scheme as **c** and with the y-axis aligned with **c. e)** Odds ratios (y-axis) when comparing the number of cases of each PRS (different panels) across the deciles (x-axis) with the middle quintile (40-60% of the distribution).

Since some of the 14 PRS target the same or similar cancer types and others only have a relatively low predictive power, we selected the top 8 different PRS that predict 8 different cancer types (breast, colorectal, prostate, glioma, thyroid, endometrial, melanoma and kidney) to continue our analyses (**Figure 1c**). Moreover, when including each these 8 PGRS to a simple model with demographic variables (age and sex), the accuracy of the model increased, showing that the predictive power of PGRS is independent of these demographic variables (**Figure S4**).

Finally, we also tested to what extent the PGRS built for a specific cancer could also be predicting other forms of the disease. To test that idea, we extended the fold-enrichment analysis per deciles to see if other cancer types than the intended one also showed an enrichment at one of the extremes (**Figure S5**). By that measure, four of the eight PGRS that we studied (breast, colorectal, kidney and melanoma) are, indeed, specific to the intended cancer type. On the other hand, the remaining PGRS (glioma, thyroid, endometrial and prostate) where depleted in cases of other cancer types at the top 20% of individuals, suggesting a negative genetic correlation between the target cancer type and other forms of the disease. Remarkably, the cross-predictions between the PGRSs and other cancer types were not consistent with each other. For example, while there were less endometrial cancer patients amongst those with the highest thyroid cancer PGRS, the opposite is not true, as the number of thyroid cancer patients at the top of the endometrial cancer PGRS is approximately the expect (OR ~ 0.9, p > 0.7). Moreover, there are no correlations between the PGRS of cancer types that cross-predict (**Figure S1**).

### Interactions between cancer subtypes and PRS

Some cancer PRS are known to predict with better accuracy specific subtypes of the disease. For example, the PRS for breast cancer has better predictive power for those tumors that are estrogen receptor positive rather than those that are estrogen receptor negative^5^. Such preferential predictive power points to interactions between the germline and somatic genomes allowing us, among others, to improve the predictive power of PRS and better understand the oncogenic mechanisms of these genetic variants. Therefore, quantifying and understanding the interactions between PRS and the landscape of somatic variants in different cancer types is of utmost importance. We used the deep molecular characterization of tumor samples in TCGA to explore this phenomenon with deeper detail.

We first tested whether the association between PRS and different subtypes in the 8 cancer types that we could predict with a PRS. Overall, we detected statistically significant differences in the PRS values of molecular subtypes of prostate (p < 0.01), thyroid (p < 2e-4) and endometrial cancer (p < 0.001) (**Figure 2**). In prostate cancer, we found marked differences between the two most predominant subtypes in TCGA, “ERG” and “other”, with the latter having much higher PRS than the former. The “Other” subtype includes all samples without mutations in ERG, ETV1, ETV4, FLI1, SPOP, IDH1 and FOXA1. These are samples with mutations in PTEN, TP53 or PIK3CA among others (**Figure 2a-d**). In endometrial cancer the subtype copy-number low was the one with the highest PRS values, whereas samples from the MSI subtype had the same PRS values as samples from other cancer types and patients with the copy number high subtype had intermediate PRS (**Figure 2e-h)**. In the case of thyroid cancer, the samples that belong to the mRNA subtype 5 had the highest PRS values, whereas the samples belonging to the other subtypes had PRS values only slightly above averages (**Figure 2i-l**).

**Figure 2.**
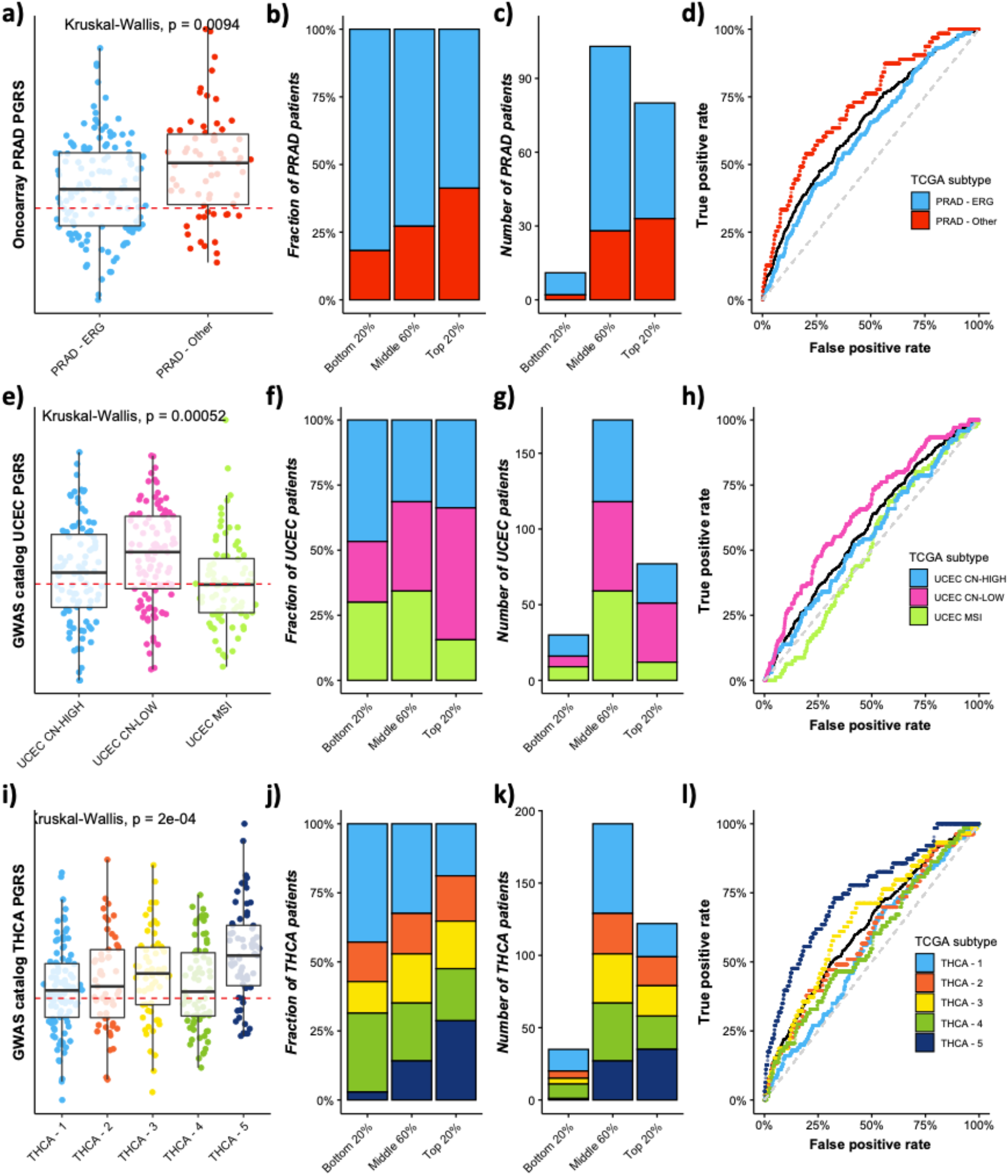
Polygenic risk scores can be biased towards specific cancer molecular subtypes. **a)** Boxplot showing the distribution of PRS (y-axis) depending on the prostate cancer subtype (x-axis). **b)** Barplot showing the fraction of samples (y-axis) that belong to the different prostate cancer molecular subtypes depending on the PRS values (x-axis). **c)** Same as **b** but in absolute value (y-axis). **d)** ROC curves for the different molecular subtypes. Curves are colored depending on the prostate cancer subtype. The original prostate cancer curve is shown in black and the diagonal as a trimmed gray line. Plots **e-h** show the same data but for endometrial cancer and plots **i-l** for thyroid cancer.

### Patients with tumors driven by certain somatic driver mutations have different PRS

The prevalence of some somatic driver mutations correlates with the ancestry of cancer patients^22^ or the germline alleles in certain loci^13^. This, together with the differences in the PRS values of tumors belonging to different somatic molecular subtypes (**Figure 2**), made us wonder if our PRS could also correlate with the presence of somatic driver mutations.

To test this hypothesis, we compared the PRS values of tumors with and without the most common driver mutations in each cancer type (those present in at least 20 patients). Our analysis revealed four different associations between PRS and somatic driver mutations (FDR < 0.1, **Figure S6**). In breast cancer, samples with TP53 somatic mutations have lower PRS values. Given the higher prevalence of TP53 mutations in ER negative breast tumors, this agrees with this breast cancer subtype having higher PRS than ER positive tumors^5^. In the case of thyroid papillary carcinoma, tumors with mutations in BRAF have higher PRS values, agreeing with the molecular subtype association that we found (**Figure 3**). We also found that brain tumors with somatic mutations in TGFBR2 and PTEN had lower and higher PRS values respectively (**Figure S6**). Finally, since both, the frequency of some driver alterations^23^ and the effect of some germline risk variants^24^, can have different effects depending on the biological sex, we thought that it is likely that biological sex also plays a role in the interactions between PRS and the somatic landscape of the tumors. To test this hypothesis, we tested whether the association between PRS and driver events was different depending on the biological sex (**Figure S7**). We only found two such instances, (FDR < 0.2): APC in colorectal cancer and CDKN2A in kidney clear cell carcinoma. In the first case the PRS values of female colorectal cancer patients are higher if their tumor also has an APC mutation than if their tumor has not. This difference is not seen in male patients. In the second example, female kidney cancer patients have higher PRS if their tumors also have a CDKN2A somatic mutation, whereas this is not the case in male patients. Overall, our results show potential interactions between PRS and somatic driver events.

**Figure 3.**
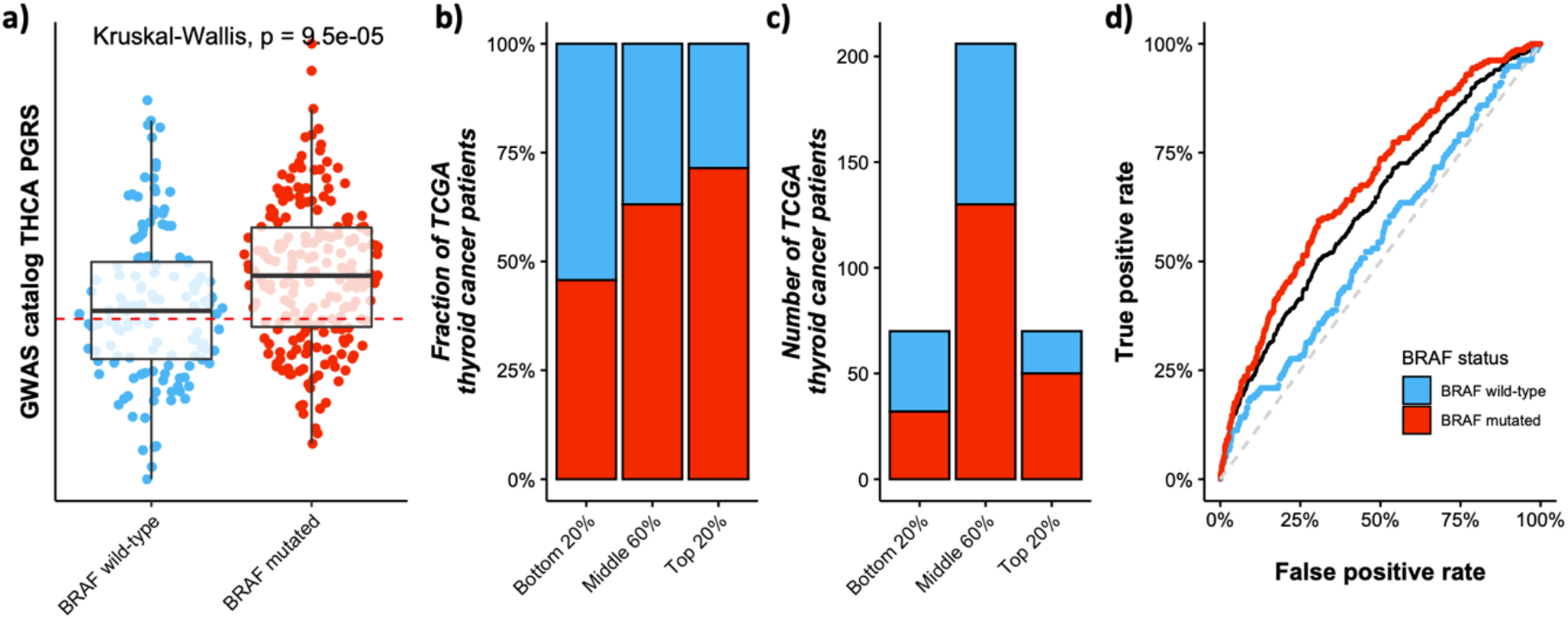
Patients with thyroid tumors driven by BRAF somatic mutations have higher PRS. **a)** Boxplot showing the distribution of PRS (y-axis) depending on the somatic mutation status of the thyroid tumor (x-axis). **b)** Barplot showing the fraction of samples (y-axis) that are BRAF mutated (red) or BRAF wild-type (blue) on the PRS values (x-axis). **c)** Same as **b** but in absolute value (y-axis). **d)** ROC curves for the thyroid tumor depending on their BRAF status. The original thyroid cancer curve is shown in black and the diagonal as a trimmed gray line.

### Towards driver-specific PRS

So far, we have described correlations between PRS and different aspects of the somatic evolution of the tumor, such as the molecular subtype or the presence of certain somatic driver mutations. However, since each PRS summarizes the risk across all loci known to be associated with a certain cancer type, it is possible that these somatic events only correlate with a subset of all the loci that make a PRS. In that case, the correlation between the germline genome and the somatic events would be obscured as the PRS includes loci that do not correlate with the somatic event of interest.

For example, our brain cancer PRS does not distinguish between IDH1 mutated and IDH1 wild-type brain cancers (**Figure 4, left**). However, Eckel-Passow *et. al.* recently identified a subset of 8 glioma risk loci that correlate with somatic IDH1 mutations^25^. Following their findings, we used their classification to deconvolute our brain cancer PRS into two different ones: one PRS specific to IDH1-mutated gliomas and another for non-IDH1 mutated gliomas. Reproducing the findings from Eckel-Passow *et. al.*, our brain cancer IDH1-mutated and the IDH1 wild-type PRS were specific for those tumors with or without an IDH1 somatic mutation (**Figure 4**, middle and right, respectively). Notably, the IDH1-mutated PRS was specific for brain cancers, as patients with somatic IDH1 mutations from other cancer types had the same PRS values as those with no IDH1 mutations. These results suggest that it might be possible to deconvolute the existing PRS for different cancer types to build driver-specific PRS that predict certain cancer-type and driver event combinations.

**Figure 4.**
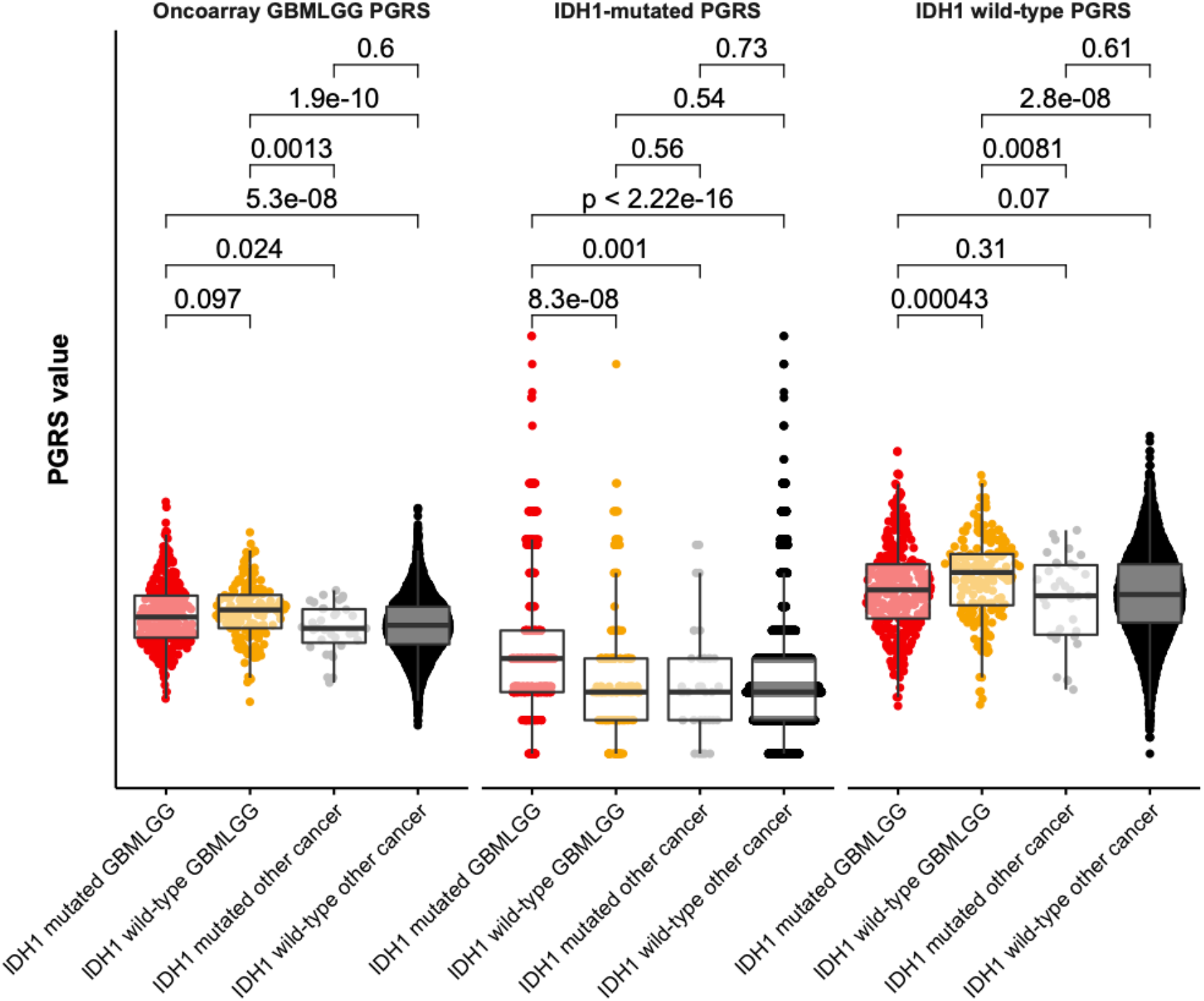
Deconvolution of glioma PRS. We classified the germline variants from the original glioma PRS depending on whether they are associated with IDH1 somatic mutations or not to create two new PRS. Then, we calculated the values of these three PRS (glioma -left panel-, IDH1-mutated glioma -middle panel- or IDH1 wild-type glioma -right panel-) for each TCGA patient (y-axis). Each dot represents a TCGA patient, and they are grouped according to their IDH1 somatic mutation status and whether they are glioma/glioblastoma patients or not.

To try to build driver-specific PRS with TCGA data, we looked for associations between the individual germline variants from each PRS and the different somatic driver genes in each cancer type. However, our overall results show that the limited sample size of TCGA does not seem to have enough statistical power to find enough associations between germline variants and somatic driver genes to build other driver-specific PRS (**Figure S7**). This, together with the associations that we previously identified between PRS and cancer subtypes and driver genes, highlight the importance of generating cohorts with matching germline and somatic mutation profiles.

## Discussion

Polygenic scores (PRS) can potentially help identify patients with different genetic risk of suffering many diseases^26^, including various forms of cancer. However, given the growing evidence showing widespread associations between the germline and somatic genomes in cancer, we need to better understand how they associate with other important variables, such as tumor subtypes^5^ or other somatic alterations^25^.

Here we used data from The Cancer Genome Atlas (TCGA) to better understand how PRS for different cancer types might preferentially predict patients with tumors that belong to certain subtypes or have specific somatic mutations. First, we validated some PRS in TCGA. Out of the 27 different PRS that we tested, 16 predicted the correct cancer type (AUC > 0.55 and Fisher’s test enrichment in cases between extreme quintiles FDR < 0.05). Since TCGA has a thorough characterization of the somatic genome, this allowed us to find associations between eight of these PRS and tumor subtypes or somatic cancer driver events. Finally, we also found evidence that other associations between PRS and somatic alterations might yet still to be discovered, but we could not detect them likely due to relatively low statistical power of TCGA.

There are different potential explanations for the association of these PRS with specific tumor subtypes. For example, some tumor subtypes are more common than others, which could bias the variants and weights identified in GWAS. This would, in turn, affect the composition and predictive power of the PRS derived from the GWAS results. This could be the case of the bias that we observed in prostate cancer subtypes. While there aren’t yet comprehensive molecular epidemiological studies looking at the prevalence of somatic alterations in different cancer types, ERG-driven prostate tumors represent around a quarter of the 1627 tumors analyzed in the ACCR-GENIE^27^ database (418 tumors, 25%), whereas almost half (788 tumors, 48%) belong the “other” subtype. If the composition of the prostate cancer GWAS studies follows this proportion, then the variants identified would likely be biased towards the “other” subtype, making it more predictable than the ERG-driven tumors.

Another possibility is that some cancer subtypes might depend more on the germline background, whereas others might be more driven by the somatic mutations their cells acquire. This could potentially be the case of the endometrial cancer associations we observed, where patients with tumors from the copy-number low subtype have higher PRS than the ones with tumors that are either copy-number high or have microsatellite instability (MSI). Our hypothesis is that, after a certain number of somatic alterations, a cell can become cancerous regardless of its germline background. This hypothesis agrees with our observation that MSI and copy-number high endometrial tumors can barely be predicted with the endometrial cancer PRS. On the other hand, if an individual has a pro-oncogenic germline background, a few somatic alterations might be enough to trigger cancer growth. This would be the case of the copy-number low subtype.

A third possibility is that the genetic architecture of different cancer subtypes is different. This could be the case of the association we found between thyroid cancer PRS and BRAF somatic mutations. In this case it could be that acquiring and BRAF mutation is more likely to end in thyroid cancer depending on the germline background of the patient, particularly the alleles at rs1588635. Moreover, our analysis grouped together all the different somatic alterations in a given driver gene. We and others have previously shown that different mutations within the same gene can lead to distinct phenotypes. The same is true for different types of alterations: a missense mutation in a gene might have a different effect than a copy-number alteration or a methylation. Therefore, it is also possible that each somatic mutation and/or mutation type within a driver gene depends or interacts differently with the germline background of the patient.

Finally, we also showed how PRS can potentially be deconvoluted to identify subsets of variants that create new driver-specific PRS. In our case we validated the brain cancer IDH1-mutated and a brain cancer IDH1 wild-type PRS by classifying the variants from the glioma PRS according to their association with IDH1 mutation status. While TCGA does not have enough statistical power to deconvolute the rest of PRS, we believe that bigger cohorts with matching germline and somatic data should be able to identify new associations that allow to build more driver-specific PRS. These could, potentially, be combined with other tests that look for somatic mutations in cell-free DNA^28^ to diagnose suspected cancer patients. We expect that the combination of both approaches would outperform each test alone.

## Clinical implications

The clinical usefulness of a PRS is largely determined by its predictive power (albeit other factors such as disease incidence, the type of available interventions for the disease or the intended use of the PRS also play an important role). For instance, a recent study^29^ suggested that a breast cancer PRS can help decide mammography screening strategies even if it has a modest AUC (0.6-0.7), but can only be useful for more radical clinical interventions that can have serious adverse effects, such as preventive use of tamoxifen, once it reaches a much higher discriminatory power (AUC > 0.8). It is clear that none of the PRS that we studied have this predictive power at the cancer type level. However, our molecular subtype analysis revealed that, in some cases, the AUC of the subtypes that are better predicted is notably higher than that of the global cancer type. For example, in prostate cancer the PRS has an overall AUC of 0.65 and the odds ratio of cases between the extreme quintiles is 5.5-fold (p < 1e-15). For tumors from the subtype “other” these values increase to 0.72 and 16-fold (p < 7e-8) respectively. In endometrial cancer patients, the original AUC is 0.59 and the enrichment in cases is 2.5-fold (p < 2e-6). For the copy-number low subtype these numbers increase to 0.65 and 5.8-fold (p < 2e-6) respectively. In the case of thyroid cancer, the original values are 0.63 for the AUC and 3.8-fold enrichment (p < 5e-14), which increase to 0.75 and 35 fold-enrichment (p < 8e-10) for the THCA-5 subtype. These values are much closer to being clinically useful, even more when considering the BRAF-mutated thyroid cancer is the most aggressive form of the disease^29^.

Our study also has several limitations. For example, since the predictive power of PRS can depend on the genetic ancestry of the individual^30^, we focused our analyses on TCGA patients with European ancestry. Therefore, we do not know to what extent our results can be extrapolated to other ancestries. However, the frequency of somatic alterations in some cancer genes varies depending on the genetic ancestry of the patient^22^, so it is likely that PRS built with data from other ancestries can also have similar biases as to the ones we have observed. Also, TCGA is relatively small, making it unfit to comprehensively explore the landscape of interactions between PRS and somatic properties of tumors, particularly for driver events, as most of them are only present in a few dozens of patients.

In conclusion, our results show that the accuracy of PRS for any cancer type can differ depending on the different somatic properties of the tumor. Together with recent evidence that PRS predictive power differs depending on sex, age or socio-economic status^31^, our results suggest that should PRS be widely adopted in screening and cancer prevention programs, this could have important consequences in which subtypes are properly diagnosed. Moreover, since GWAS studies have mostly been done without accounting for the somatic subtypes and driver alterations, it is likely that the biases detected here are only a small fraction of the total. We believe that creating cohorts with matching germline and somatic genomes will identify new germline-somatic interactions and help find new germline loci that are specific of certain tumor features. This will improve the predictive power of PRS, which is critical before their widespread implementation in the clinic.

## Acknowledgements

We would like to thank all the patients and scientists involved in The Cancer Genome Atlas. We also would like to thank Urko M. Marigorta, David Tamborero and Collin Tokheim for their previous feedback about the project. E.P-P received support from a La Caixa Junior Leader Fellowship from Fundació Bancaria La Caixa. A.V received support from Institució Catalana deRecerca Avançada (ICREA). E.Z. received funding from R01CA227466 and K24CA169004. R.W.S. is supported by a postdoctoral fellowship from the National Cancer Institute (NCI) T32CA221709.

## Author contributions

Conceptualization: E.P-P, A.V, E.Z. Analysis, computation and software: E.P-P, R.W.S. Interpretation: all authors. Writing, review and editing: E.P-P, A.V. Funding Acquisition: E.P-P, A.V, E.Z.

## Methods

### Data and code availability

The TCGA birdseed genotyping data and clinical data can be found at the legacy archive of the GDC (https://portal.gdc.cancer.gov/legacy-archive). The Cancer Genome Atlas (TCGA) quality-controlled Genome Wide SNP 6.0 genotyping data imputed to Haplotype Reference Consortium are controlled access files and will be made accessible to researchers with proper dbGAP authorization. These data have been deposited to Synapse (Available upon acceptance for publication).

### Curation of the germline genotypes from The Cancer Genome Atlas

Germline genotype data for common variants used in heritability analysis and Genome-Wide Association Studies (GWAS) were obtained from Affymetrix Genome Wide SNP 6.0 arrays (TCGA legacy archive https://portal.gdc.cancer.gov/legacy-archive). Birdseed genotyping files representing 905,600 variants for 11,521 samples were downloaded.

Birdseed files were read in R v3.5.0 using the Affymetrix SNP Array 6.0 (release 35) annotation file, and 905,422 variants were successfully loaded and analyzed in PLINK version 1.9. Samples were cross-referenced against previously whitelisted genotyping samples^32^. Based on established TCGA barcode identifiers, samples annotated with Analyte code “G” (Whole Genome Amplification) were further excluded. A final set of 10,946 whitelisted samples with Analyte code “D” (DNA) were retained for quality assessment.

Stringent quality control measures were applied to the SNP genotyping data^33^. SNPs and individuals with greater than 5% missingness were excluded; leaving a total of 861,351 variants and 10,917 samples for subsequent analysis.

Initial Principal Component Analysis (PCA) ancestry analysis was performed to facilitate heterozygosity calculations. PCA without linkage-disequilibrium pruning was performed in PLINK 1.9 (Chang et al., 2015), and visual examination of the concordance of the principal component plots with the self-reported race and ethnicity annotations revealed that the first 3-4 PCs captured population structure information, while PCs 5-6 captured outliers. PCA initial ancestry clusters were determined by performing both k-means and partition around medoids (PAM) clustering on either the first three or first four PCs. We computed gap statistics and average silhouette widths iteratively for number of clusters, k=1 to 10 for k-means and PAM methods respectively to find the optimal number of clusters for each method. We found PAM using the three PCs yielding 4 optimal clusters to show high concordance with self-reported race/ethnicity (ancestry cluster 1 = European, cluster 2 = Asian, cluster 3 = African, cluster 4 = American). Based on the initial ancestry cluster assignments, heterozygosity was calculated in PLINK 1.9 within each initial PCA-based ancestry cluster and a total of 250 samples with heterozygosity >3*SD above the ancestry mean were removed.

Selection of a representative sample for each individual was then conducted. Individuals represented by more than one sample, blood-derived normal samples were preferentially selected; for those with more than one blood-derived samples, samples with higher call rates were retained. After these steps, a total of 10,128 unique individuals remained for subsequent analysis.

Final filtering steps for SNPs were conducted across the 10,128 unique individuals and restricted to autosomal chrs. Hardy-Weinberg Equilibrium (HWE) was calculated in PLINK 1.9 across individuals within largest ancestry cluster (European ancestry cluster 1). SNPs that deviated from the expectation under HWE (p < 1×10−6) within the European ancestry cluster were excluded with the exception of SNPs previously associated with any cancer as reported in the GWAS catalog (p < 5×10−8) (Rashkin et al., 2019) since they may deviate from HWE in cancer patients. Minor allele frequency (MAF) was calculated and variants with MAF < 0.005 were excluded. Finally, duplicate SNPs with identical genomic first position were removed. A total of 838,948 autosomal chr variants for 10,128 unique individuals passed after the aforementioned QC steps.

### Genotype imputation

The quality-controlled genotyping file was stranded and imputed against the Haplotype Reference Consortium (HRC) (Loh et al., 2016a; McCarthy et al., 2016). Prior to HRC stranding, all palindromic SNPs (A/T or G/C) were removed. Stranding was then performed using the McCarthy Group tools (HRC-1000G-check-bim-v4.29), which compares our data genotyping alleles to the corresponding SNP alleles from HRC (v1.1 HRC.r1-1.GRCh37.wgs.mac5.sites.tab), leaving 680,389 correctly matched variants for imputation. Phasing and imputation were performed using a standard pipeline on the Michigan Imputation Server (MIS). Phasing was performed using Eagle version v2.3 (Loh et al., 2016b) on the variant call file (VCF). To reduce the run time, the VCF file was divided into 22 files corresponding to individual autosomal chromosomes. By default, Eagle restricts analysis to bi-allelic variants that exist in both the target and reference data. Minimac3 was used to run the imputation. For each of the 22 VCF files, the MIS breaks the dataset into non-overlapping chunks prior to imputation. For HRC imputation, the HRC r1.1.2016 reference panel was selected using mixed population for QC, with a total of 39,127,678 SNPs returned after imputation.

### Final ancestry calls

PCA was performed on the final quality-controlled genotyping file and final PAM-based ancestry clusters were computed for the 10,128 individuals for optimal k=4. We found very high concordance of initial and final ancestry assignments (99.98% matching, the 2 samples varying between initial and final ancestry cluster computation assigned to NA).

The four ancestry cluster are as follows: (1) PAM ancestry cluster 1 is concordant with European ancestry, capturing 97.27% of individuals self-reporting as White, as well as 82.16% of individuals with self-reported non-Hispanic/non-Latino ancestry and 45.96% with self-reported Hispanic/Latino ancestry; (2) ancestry cluster 2 with African ancestry, capturing of 97.53% of individuals self-reporting as Black/African-American race; (3) ancestry cluster 3 with Asian ancestry, capturing 90.88% of individuals self-reporting as Asian and 88.89% self-reporting as Native Hawaiian/Pacific Islander; and (4) ancestry cluster 4 with a subgroup of individuals with American ancestry capturing 60% of individuals self-reporting as American Indian /Alaska Native and 47.2% with self-reported Hispanic/Latino ethnicity (Carrot-Zhang J et al, manuscript submitted).

PC’s 1-7 show further population sub-structure in the Asian and European ancestry clusters. PAM ancestry sub-clusters were computed using PC’s 1-7 for individuals within the Asian ancestry cluster which yielded two optimal sub-clusters, and within the European ancestry cluster which yielded three optimal sub-clusters (GDC Publication Page Figure S2-B). Of note, 72.46% of European sub-cluster 3 self-reports as Asian (15.94% have no race reported). Ancestry clusters, sub-clusters, self-reported race and ethnicity and PC’s 1-7 are provided for each individual. We used in our analyses only those samples with genetic European ancestry (n = 8304).

### Building the polygenic risk scores from the GWAS catalog

We downloaded the GWAS Catalog on May 2020. We then manually filtered the variants to include only those associated to cancer, obtained in studies from european population, with more than 2000 individuals and a p-value below 5e-8. We used liftover to map the variants from GRCh38 to GRCh37. We assigned the weights associated to each variant as the log of the odds ratio for the risk allele to the cancer type according to the GWAS catalog. Finally, we manually mapped the different cancer phenotypes to the cancer types studied in TCGA.

### Building the polygenic risk scores from Oncoarray

For the breast cancer PRS we used the 313 SNPs identified in Mavaddat *et. al.*^5^. According to their study, they first identified 305 SNPs associated with overall breast cancer (p < 1e-5). This 305-SNP PRS was supplemented with 6 additional SNPs associated with ER-positive at p value < 10−6 and, in addition, by two known rare breast cancer susceptibility variants in the BRCA2 and CHEK2 genes, bringing the total number of SNPs included to 313.

In the case of colorectal cancer we used the variants, risk alleles and odds ratios from Law *et. al.*^20^. For glioma and glioblastoma we used those described in Melin *et. al.* ^18^ and for ovarian cancer those from Phelan *et. al.* ^21^.

Finally, for the prostate cancer, lung squamous carcinoma and lung adenocarcinoma PRS we downloaded the summary data from Schumacher *et. al.* ^2^ and McKay *et. al.* ^19^, filtered all the alleles that did not reach genome-wide significance (p > 5e-8) and used plink to clump all the remaining variants with the following parameters: --clump-p1 5e-8 --clump-r2 0.1

### Evaluation of the PRS

We calculated the PRS for each patient for each cancer type with plink using the --score parameter. The correlation between PRS for different cancer types was calculated with the R package “corrplot” using the “hclust” function.

The fold-enrichment analyses were calculated using a two-sided Fisher’s test. We compared the number of patients from the intended cancer type in different deciles of TCGA (10%, 20%, 30%, 70%, 80% and 90%) to those between the 40% and 60% of the distribution.

To calculate the AUC of the different PRS we used the R package “pROC” and set the patients from the intended cancer type as positives and the rest of TCGA patients as negatives. For sex-specific cancers (prostate, endometrial, testicular and ovarian) as well as for breast cancer we used only the subset of TCGA with the appropriate sex for our analyses.

To compare the performance of the PRS to the model with demographic variables (age and, when appropriate, sex), we created three models: one with only the PRS values, another with the demographic variables and a third with both. Then, for each model, we calculated the Nagelkerke’s pseudo-R and the AUC with R packages “rcompanion” and “pROC” respectively.

Finally, to calculate the cross-cancer predictions we used the same approach as for the fold-enrichment analysis explained above, but using each of the not-intended cancer types as the target cancer for each PRS.

### Correlation between PRS and cancer subtypes

We used the cancer subtypes from TCGAbiolinks^34^, available in Thorsson *et. al.*^32^. The subtypes are defined according to the main subtype used in the TCGA publication describing each cohort from each cancer type. Then, for each of the final 8 PRS that we analyzed in detail, we selected subtypes with 50 or more patients of European Ancestry in the intended cancer type. Finally, we compared the PRS in each cancer subtype using the Kruskal-Wallis test from the R package “ggpubr”.

### Correlation between PRS and driver events

We used the cancer driver events from Sanchez-Vega *et. al.*^35^. In brief, they considered a cancer driver gene as somatically mutated if it was affected by a somatic mutation, copy-number variant event or change in methylation. Of the three driver levels (pathways, genes and individual variants), we used the gene-level for our analysis, but the results were similar using the pathway and individual variant levels. We also included the IDH1 driver mutations from Bailey *et. al.* ^36^, as they were not in^35^. In the end we had matching driver and PRS data for 6941 patients. We used Wilcoxon-test to compare the PRS values of the patients from the different cancer types with the somatic driver mutation to those without it.

