## Supplementary Figures for "The landscape of interactions between cancer polygenic risk scores and somatic alterations in cancer cells"

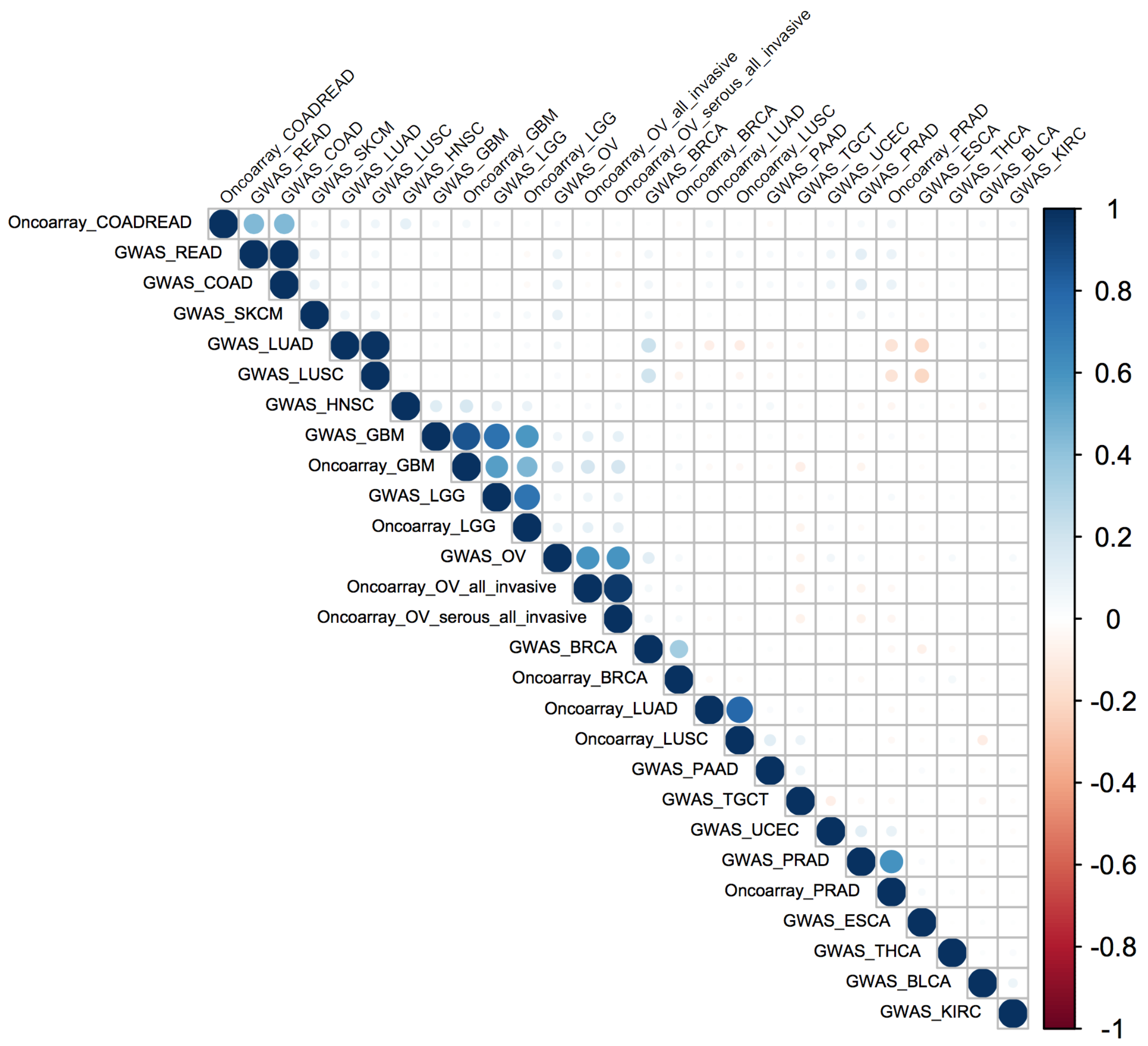


**Figure S1 - Correlation between the different PGRS in the TCGA cohort.** Each dot is colored according to the R2 value between two PGRS in blue if the correlation is positive and in red if it is negative. The correlation is calculated based on the PGRS values of all individuals in TCGA.

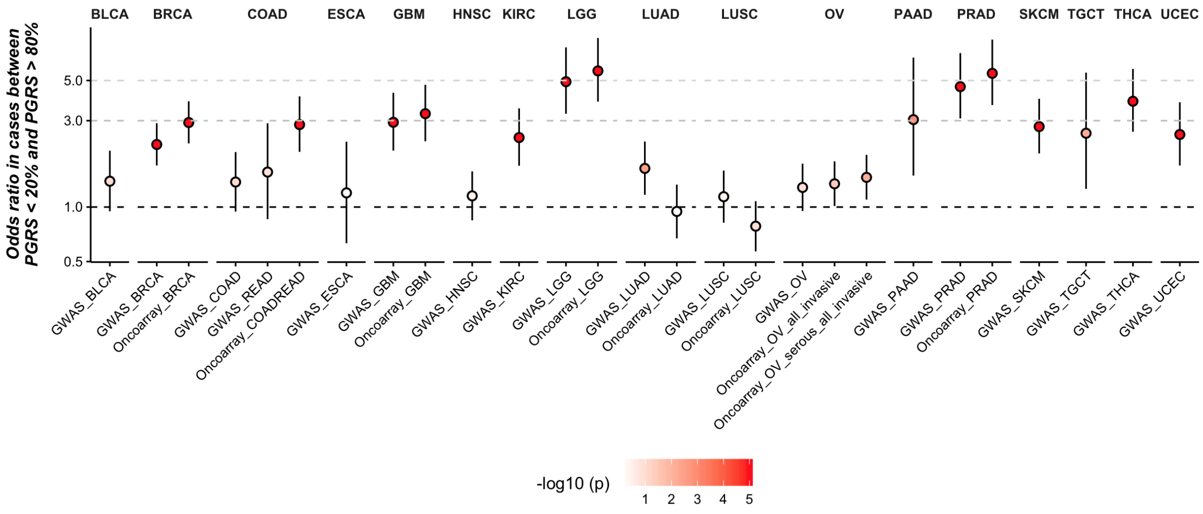


**Figure S2 - Odds ratio of the PGRS in TCGA.** Each dot represents the odds ratio between cases in TCGA individuals with a PGRS above 80% and those with a PGRS below 20%. Dots are colored according to the p-value of the Fisher test from white (p = 1) to red (p < 1e-5). Bars represent the 95% confidence interval for the odds ratio.

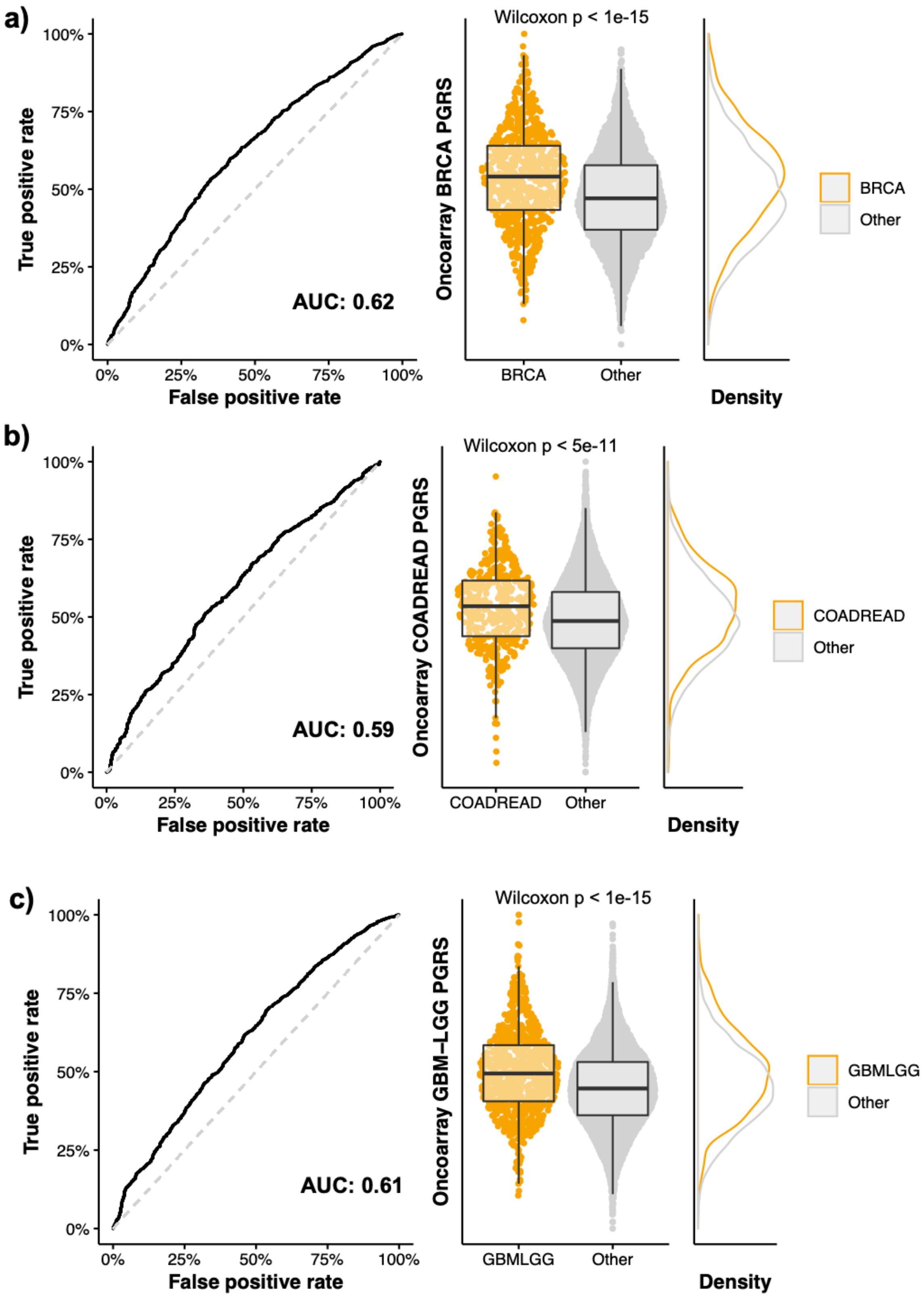


(Continues next page)

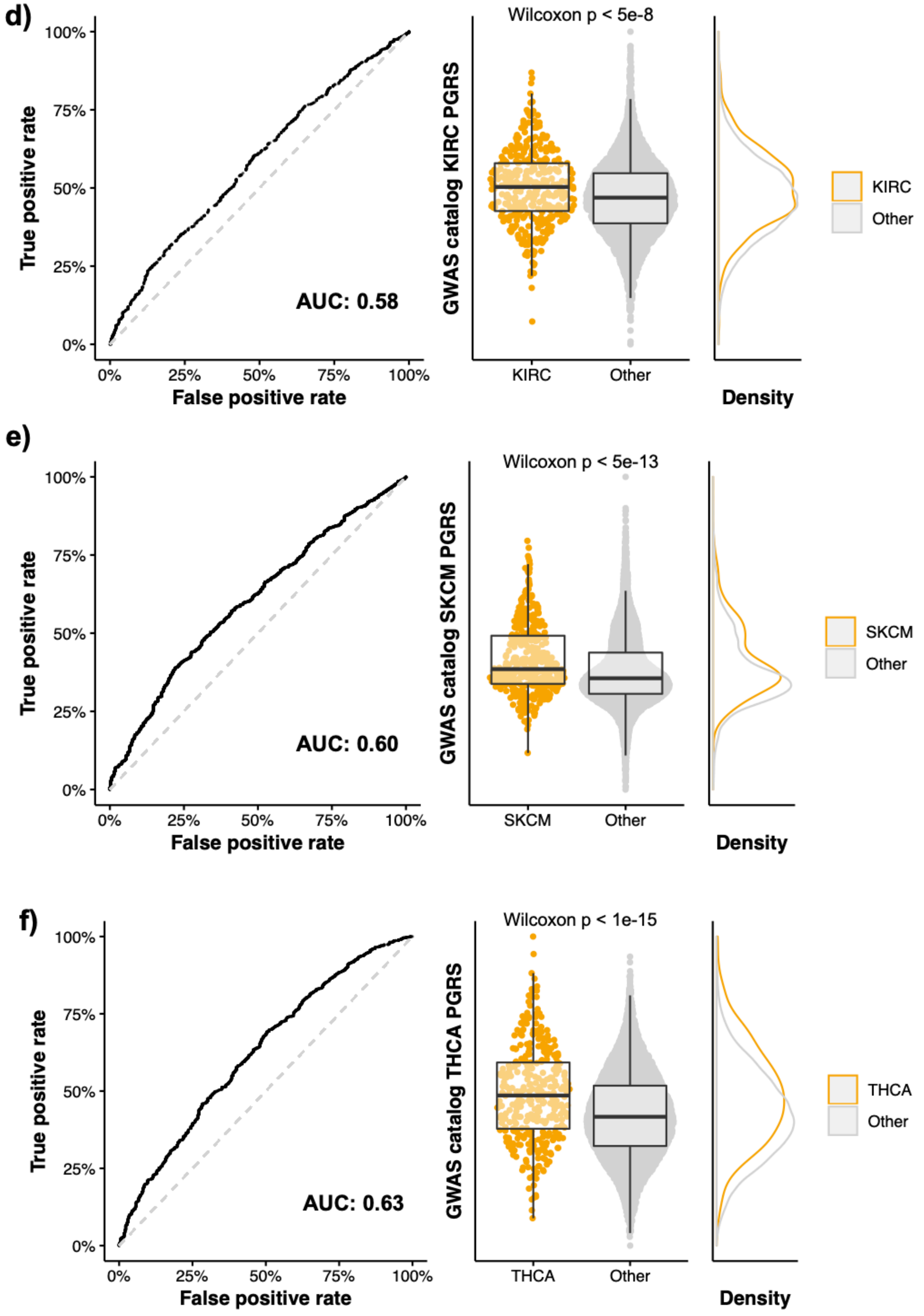


(Continues next page)

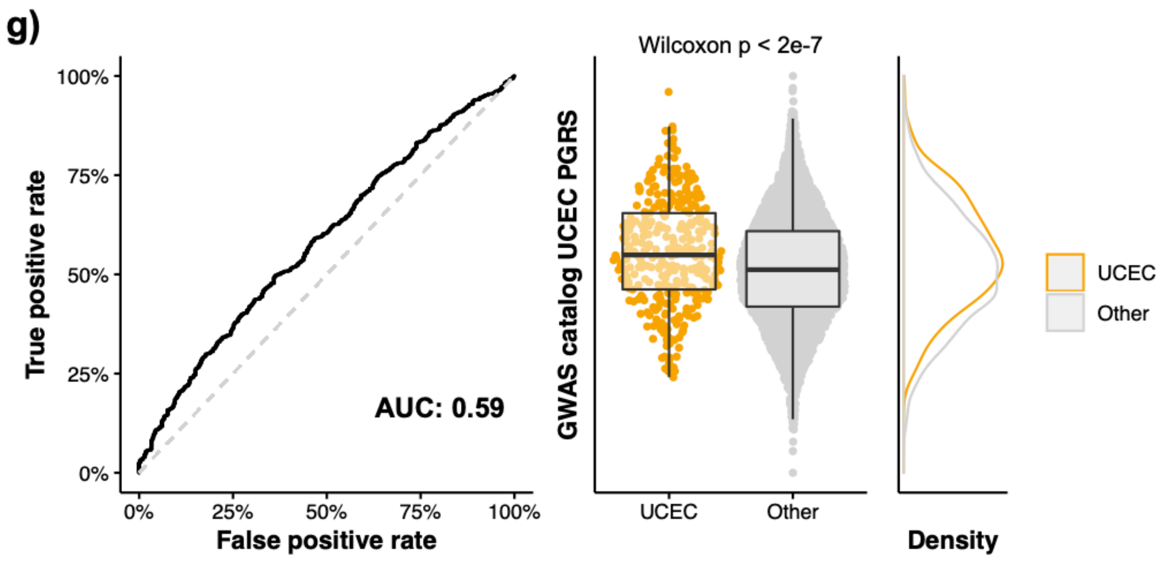


**Figure S3 – Predictive power of the selected PGRS.** We show the performance of seven of the selected PGRS: breast (a), colorectal (b), brain (c), kidney clear cell (d), melanoma (e), thyroid (f) and endometrium (g) cancers. Plots for prostate cancer are shown in Figure 2. The left plots show the ROC curve of each PGRS. The y-axis is the true positive rate and the x-axis is the false positive rate. The ROC curve is shown in black and the diagonal as a trimmed gray line. The middle plots show the difference in the distribution of PGRS values in the target cancer type (in orange) and the rest of samples in TCGA (in gray) as well as the p value of the Wilcoxon test comparing both distributions. Each dot is a TCGA sample. The right plots show the density distribution of the PGRS for the target cancer type (orange) as well as for the rest of TCGA (gray). The middle and right plots are aligned so that the density plot matches the boxplot.

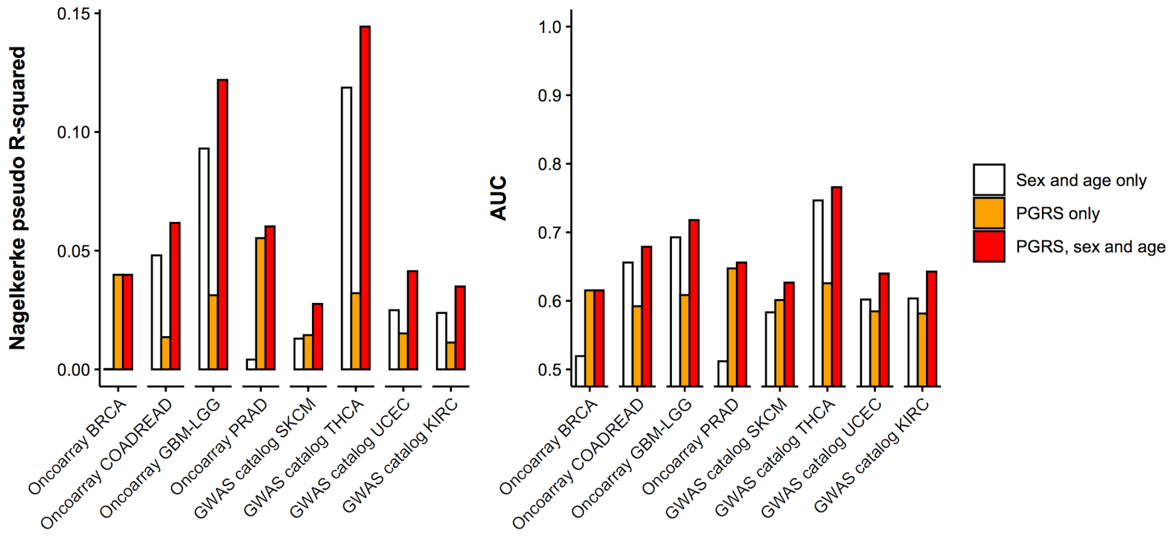


**Figure S4 - Comparison of the predictive power of demographic variables variables and the PGRS.** Each bar shows either the Nagelkerke pseudo-R2 (top) or the area under the ROC curve (bottom) when predicting the different cancer types for either demographic variables only (age and sex, white bars), the PGRS (orange bars) or both things combined (red bars). Note that sex is not included in the model for sex-specific tumors: prostate, breast and endometrium.

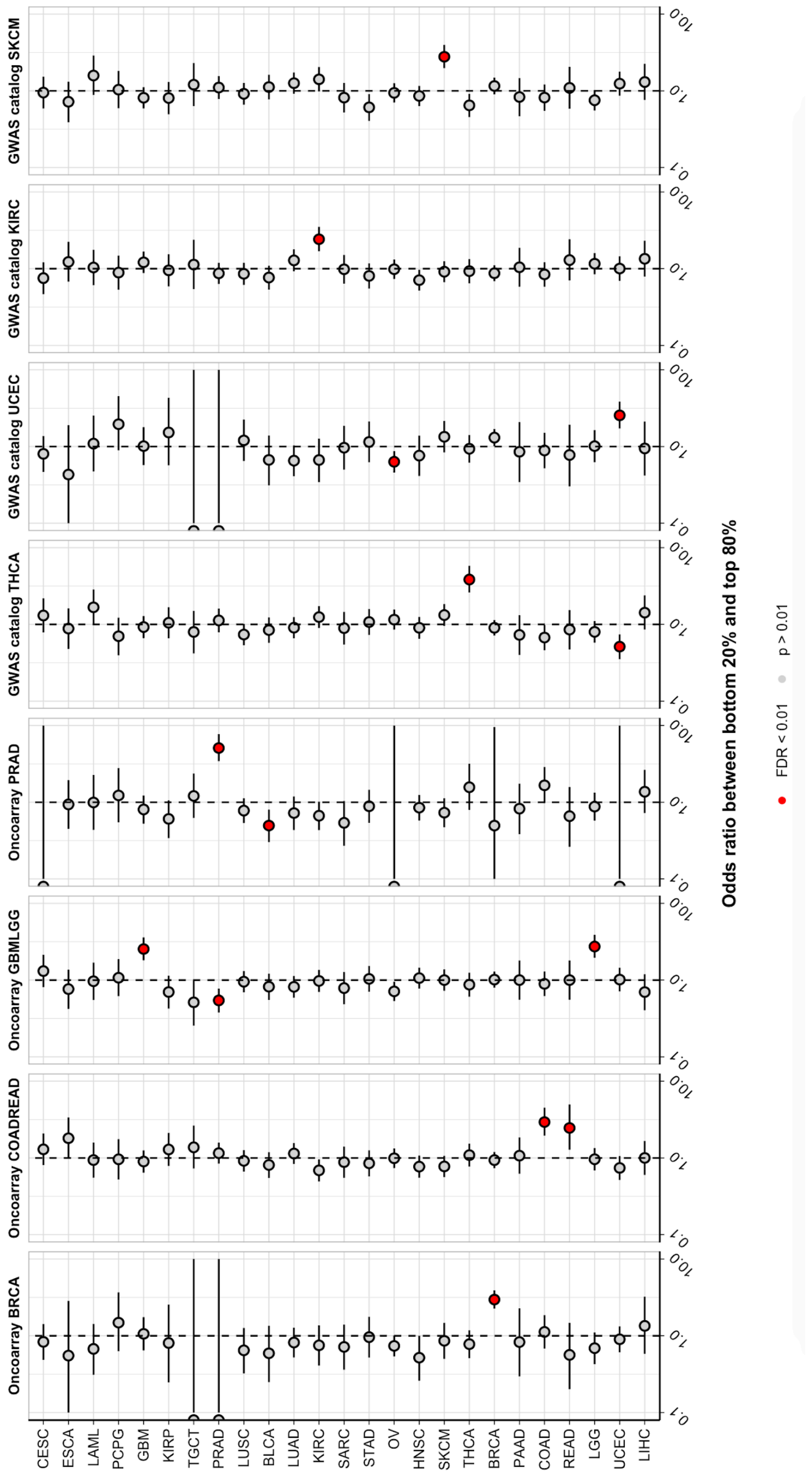


**Figure S5 - Cross-cancer predictions of the PGRS.** Each dot represents the odds ratio in cases of the different cancer types (y-axis) when comparing the top and bottom quintiles of the different PGRS. Bars represent the 95% confidence interval. Dots are colored red if the p value is below 0.01 and gray otherwise.

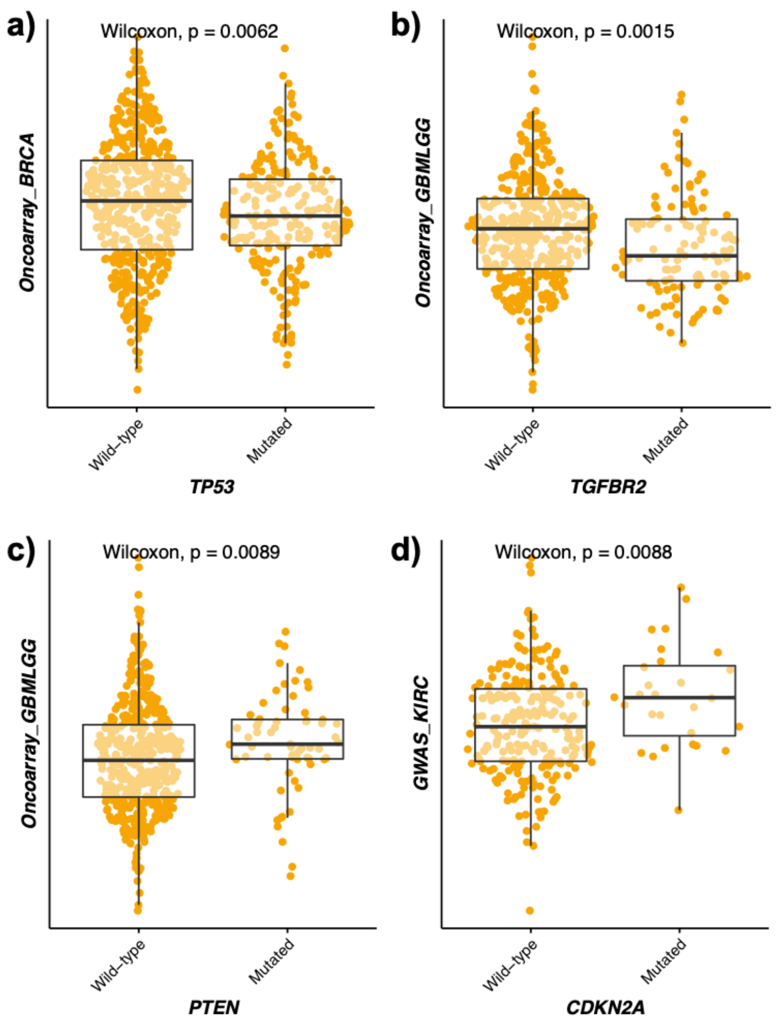


**Figure S6 – Somatic driver events that correlate with PGRS.** Each dot represents a tumor sample. Boxplots show the difference in the distribution of PGRS values (y-axis) of tumor samples (represented as dots) according to whether they have a mutation in the driver gene (x-axis). **a)** Boxplots for breast cancer samples and TP53 somatic mutations. **b)** Brain cancers and TGFBR2 somatic mutations. **c)** Brain cancers and PTEN somatic mutations. **d)** Kidney clear cell tumors and CDKN2A somatic mutations.

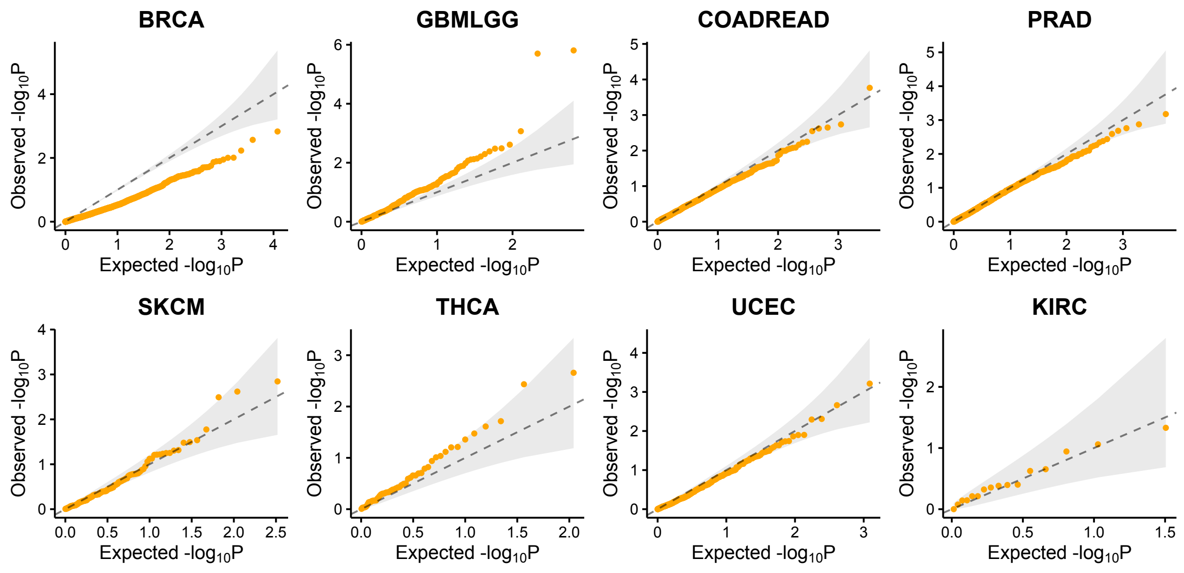


**Figure S7 – qqplots for the association tests between germline risk variants and somatic driver events.** Each plot title shows that cancer type analyzed in each qqplot, which show the observed p-values (y-axis) and the expected p-values (x-axis) distributions when doing an association analysis between germline and somatic variants in the eight cancer types of interest.
